# Meta-analysis of (single-cell method) benchmarks reveals the need for extensibility and interoperability

**DOI:** 10.1101/2022.09.22.508982

**Authors:** Anthony Sonrel, Almut Luetge, Charlotte Soneson, Izaskun Mallona, Pierre-Luc Germain, Sergey Knyazev, Jeroen Gilis, Reto Gerber, Ruth Seurinck, Dominique Paul, Emanuel Sonder, Helena L. Crowell, Imran Fanaswala, Ahmad Al-Ajami, Elyas Heidari, Stephan Schmeing, Stefan Milosavljevic, Yvan Saeys, Serghei Mangul, Mark D. Robinson

**Author notes:** these authors contributed equally and reserve the right to list themselves as first author (Anthony Sonrel, Almut Luetge, Charlotte Soneson, Izaskun Mallona, Pierre-Luc Germain).

## Abstract

Computational methods represent the lifeblood of modern molecular biology. Benchmarking is important for all methods, but with a focus here on computational methods, benchmarking is critical to dissect important steps of analysis pipelines, formally assess performance across common situations as well as edge cases, and ultimately guide users on what tools to use. Benchmarking can also be important for community building and advancing methods in a principled way. We conducted a meta-analysis of recent single-cell benchmarks to summarize the scope, extensibility, neutrality, as well as technical features and whether best practices in open data and reproducible research were followed. The results highlight that while benchmarks often make code available and are in principle reproducible, they remain difficult to extend, for example, as new methods and new ways to assess methods emerge. In addition, embracing containerization and workflow systems would enhance reusability of intermediate benchmarking results, thus also driving wider adoption.

## Introduction

Given the rapid development and uptake of new technologies in biology (e.g., high-throughput DNA sequencing, single-cell assays and imaging technologies), methodologists are presented with nearly unlimited opportunities to apply or develop computational tools to process, model, and interpret large-scale datasets across a wide range of applications. Unsurprisingly, the data explosion (Schatz 2015; Svensson, da Veiga Beltrame, and Pachter 2020) is mirrored by a massive increase in the number of computational methods; for example, at time of writing, 1,318 are listed in a database of tools for the analysis of single-cell RNA-seq data (Zappia, Phipson, and Oshlack 2018; Zappia and Theis 2021) and more than 370 tools are listed for the analysis of spatial omics data (Moses and Pachter 2022). This imposes challenges for determining what tools to use for discovery (Dance 2022). In particular, researchers often need to convince themselves that they develop or use performant tools in their research, and a typical approach is via formal benchmarks. Benchmarks can be decomposed into five steps: 1) formulating a computational task (or subtask) that will be investigated (e.g., calling differentially expressed genes); 2) collecting reference datasets by either generating (realistic) synthetic datasets or using ground truth derived from experimental data; 3) defining performance criteria (e.g., sensitivity, specificity); 4) evaluating a representative set of methods via a set of performance criteria across multiple reference datasets; and, 5) formulating conclusions and guidelines.

In terms of developing and disseminating new methods, the minimum requirement of quality is usually that the new approach provides a benefit against existing approaches. However, the current standard allows developers to be their own “judge, jury and executioner” (Norel, Rice, and Stolovitzky 2011), giving them some freedom to choose the settings and the evaluations used in the benchmark. There is a notable tension here, since methodologists that want to develop high-quality novel approaches may also be under pressure to publish (Grimes, Bauch, and Ioannidis 2018). One can argue that the risk of biases and over-optimistic evaluations of new approaches is minimised by the standard scientific review process, however this is also known to have its challenges (Tennant and Ross-Hellauer 2020). Ultimately, it is almost a foregone conclusion that a newly proposed method will report comparatively strong performance (Buchka et al. 2021; Norel, Rice, and Stolovitzky 2011). Thus, claims from individual method development papers need to be scrutinized, preferably from a neutral (i.e., independent) standpoint (Mangul, Martin, et al. 2019; Weber et al. 2019). Neutral benchmarking appears to be a popular approach, since over 60 benchmarks have been conducted for single-cell data analysis alone (see below and Additional file 1). But even if done neutrally, the community may still want a mechanism to challenge, extend or personalize the assessments (e.g., update/add reference datasets, run methods with alternative parameters, use different metrics, rank methods differently).

Neutral benchmarking shares common ground with community “challenges” for consolidating the state-of-the-art. In some subfields of computational biology, there is a long history of such challenges, such as the biannual CASP (Critical Assessment of Structure Prediction) (Kryshtafovych et al. 2021) and DREAM (Dialogue on Reverse Engineering Assessment and Methods) challenges (Stolovitzky, Monroe, and Califano 2007), where participants are invited to propose solutions to a predefined problem. The challenge model is growing in success, including leaderboards that give real-time feedback, but these can sometimes reinforce a narrow view of performance (e.g., a single measure of performance on a single dataset ("Open Problems - Multimodal Single-Cell Integration" n.d.). In addition, for a challenge to be conceptualised in the first place, the community needs to formalize and frame existing problems, have access to suitable reference datasets, gather indications that the challenge can be solved and only then assess if current technologies and methods would be able to solve it. This necessarily requires some common ground and knowledge in the field, which is often gained by neutral benchmarking studies. Although community challenges engage communities and innovation, they are typically time-gated in their scope, whereas one could also imagine benchmarking as a continuous process where challenges are integrated into the subfield’s trajectory. Initiatives in this direction include OpenEBench (Capella-Gutierrez et al. 2017), which provides a computing platform and infrastructure for benchmarking events, and ‘Open Problems in Single Cell Analysis’, which is focused on formalizing (single-cell data analysis) tasks to foster innovation in method development while providing infrastructure and datasets such that new methods can be tested (Lance et al. 2022; Luecken, Burkhardt, et al. 2022).

There are several obstacles related to running and using the results of benchmarks. In fast-moving subfields, benchmark results become rapidly out-of-date and in some cases, competing methods never get directly compared because they are developed simultaneously. Current benchmarks are always a snapshot in time, while tool development is continuous. There are common components of most benchmarks, such as reference datasets, a set of methods and metrics to score their performance, but there are typically no pervasive standards for the system or strategy of benchmarking in computational biology, except for those predefined by challenges. This lack of standards can, for example, lead to different rankings of the same methods for the same task (Luecken, Büttner, et al. 2022; Chazarra-Gil et al. 2021; Tran et al. 2020). Similarly, shortcomings of existing performance metrics may be discovered subsequent to earlier benchmarks (Lütge et al. 2021). Another area in which benchmarks underdeliver is the interpretation of performance results. Benchmark authors (or challenge organizers) generally get to make all the decisions about how the performance evaluation is conducted (e.g., how a ranking is determined, given multiple criteria).

Altogether, it remains an open question whether computational biology has reached a benchmarking optimum, achieving fair comparisons in a timely, independent and continuous manner, while also keeping the barrier low enough for colleagues, including those in adjacent fields, to participate. In this report, we review current benchmarking practices in a subfield of genome biology (single-cell data analysis) to get an understanding of where the current state-of-the-art is and based on our findings, we postulate what elements of benchmarking would be considered desirable for the future.

## State-of-the-art in benchmarking

To understand the current state of practice in benchmarking studies in computational biology, we crowdsourced a list of single-cell method benchmarks (studies were selected by the project team from the period of 2018-2021 as exhaustively as possible on the basis of having at least a preprint posted, and involving the comparison of computational methods for some form of single-cell data, and not a primary method publication; see Additional file 1 for the list of studies). We then designed a questionnaire to query various attributes of a benchmark study (see Methods, Additional file 2), and crowdsourced the review of these benchmarks. We focused on single-cell methods because it is an active area of methodological research, with an acute method explosion (Zappia, Phipson, and Oshlack 2018; Zappia and Theis 2021), and a large number of benchmarks have been conducted. The list of questions asked for each benchmark includes both factual (e.g., “Whether synthetic data is available”) and opinion (e.g., “Degree to which authors are neutral”) assessments. Overall, we queried the scope, extensibility, neutrality, open science and technical features of each benchmark. We required that at least two reviewers answer the questionnaire for each benchmark and in the case of large discrepancies between responses (disagreement in a factual question or large difference in opinion), results were consolidated manually by a third reviewer (see Additional file 1 for consolidated responses, Additional file 3 for original responses). Questions were organized into two topics: 1) overall design of the benchmark; 2) code and data availability and technical aspects.

## Overall design of benchmarks

The overall design of the 62 surveyed single-cell benchmarks, including the number of datasets, methods or criteria used for the comparison or the neutrality of the authors, were assessed. The number of benchmark datasets varied greatly, with 2 benchmarks using only 1 dataset and 1 benchmark using thousands of simulated datasets (median = 8). Likewise, the number of methods evaluated varied from 2 to 88 (median = 9) for the chosen task. Finally, the number of evaluation criteria, defined here as numerical metrics to compare method results against a ground truth, varied from 1 to 18 (median = 4), showing that current benchmarks tend to include more methods and datasets than evaluation criteria (Figure 1A). The range of analysis tasks covered by the benchmarks mirrors quite well the range of tools available (Zappia, Phipson, and Oshlack 2018; Zappia and Theis 2021). A notable exception is the category of visualisation tasks, accounting for 40% of the available tools but formally benchmarked only once (Figure 1B). 72% of the manuscripts were first released as preprints and 66% tested only default parameters (Figure 1C). We also enquired about the neutrality of the authors, defined as whether the authors of the benchmark were involved in one or several of the methods evaluated. For more than 60% of the benchmarks, the authors were completely independent of the evaluated methods (Figure 1D); neutrality is a desired attribute, although not absolutely required. More than 75% of benchmarks also assessed secondary measures (Supplementary Figure 1), such as runtime, memory usage and scalability.

**Figure 1.**
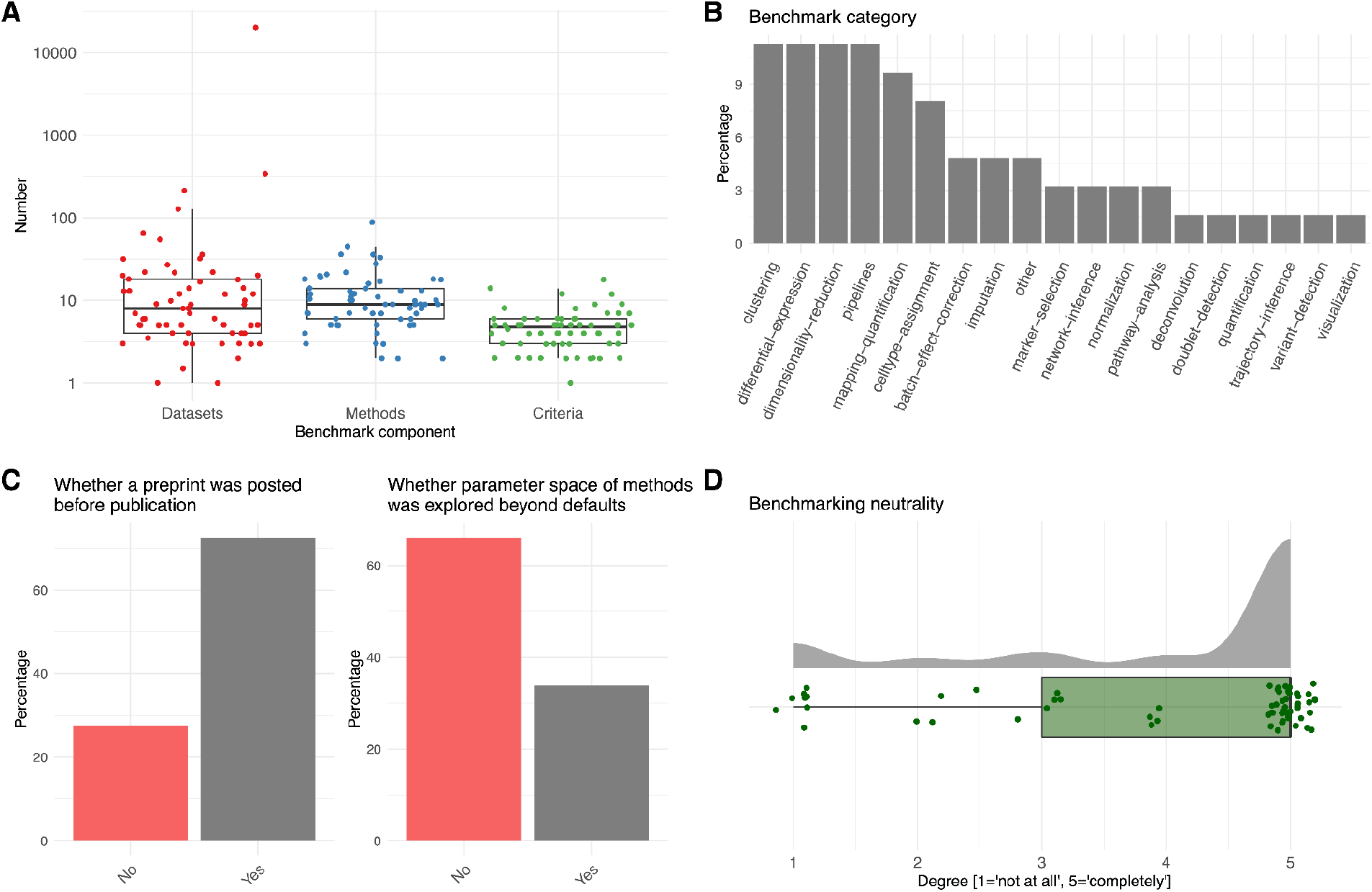
Overall design of 62 single-cell method benchmarks. Overview of crowdsourced meta-analysis across surveyed benchmarks. A) numbers of entities (datasets, methods, metrics) present in each benchmark (each dot is a benchmark). Jitter is added to the X-axis; B) data analysis tasks; C) percentages of benchmarks that were first posted as preprint or whether benchmarks explored parameter space beyond default settings; D) reviewer’s opinions on the neutrality (whether the benchmark authors were involved in methods evaluated). Jitter is added to the X-axis and Y-axis of the scores.

## Code / data availability, reproducibility and technical aspects

Also important for the uptake of benchmarking results is the open science and reproducibility practices of the studies. Thus, the second group of questions related to the availability of data, code and results as well as technical aspects. Figure 2A gives an overview of the availability of the different levels of data for benchmarking studies, highlighting that input (often ground-truth-including) data is frequently available (97% of studies). However, intermediate results, including outputs of methods run on datasets and performance results were only sparsely available at 19% and 29%, respectively. For studies that generated simulated data, less than half (19/46 articles) made their synthetic data available. Only 10% of benchmarks provided performance results in an explorable format. On the technical side, most benchmarks reported software versions of the methods being evaluated (68%), although provenance tracking (tracking of inputs, outputs, parameters, software versions, etc.) was not explicitly used. Another aspect of reproducible practice is related to workflow tools that are used to orchestrate the datasets through methods and metrics. Although their use in computational biology is increasing (Perkel 2019), we observed that less than 25% of the surveyed benchmarks used any form of them (see Figure 2B). In a similar vein, containerization of software environments is quite mature in computational biology (Gruening et al. 2018), but is rarely utilized in benchmarking (8% of studies). Concerning the methods that are compared, R and Python remain the dominant programming languages for single-cell methods, mirroring the summaries in the scrna-tools database (Zappia and Theis 2021); see also Supplementary Figure 1.

**Figure 2.**
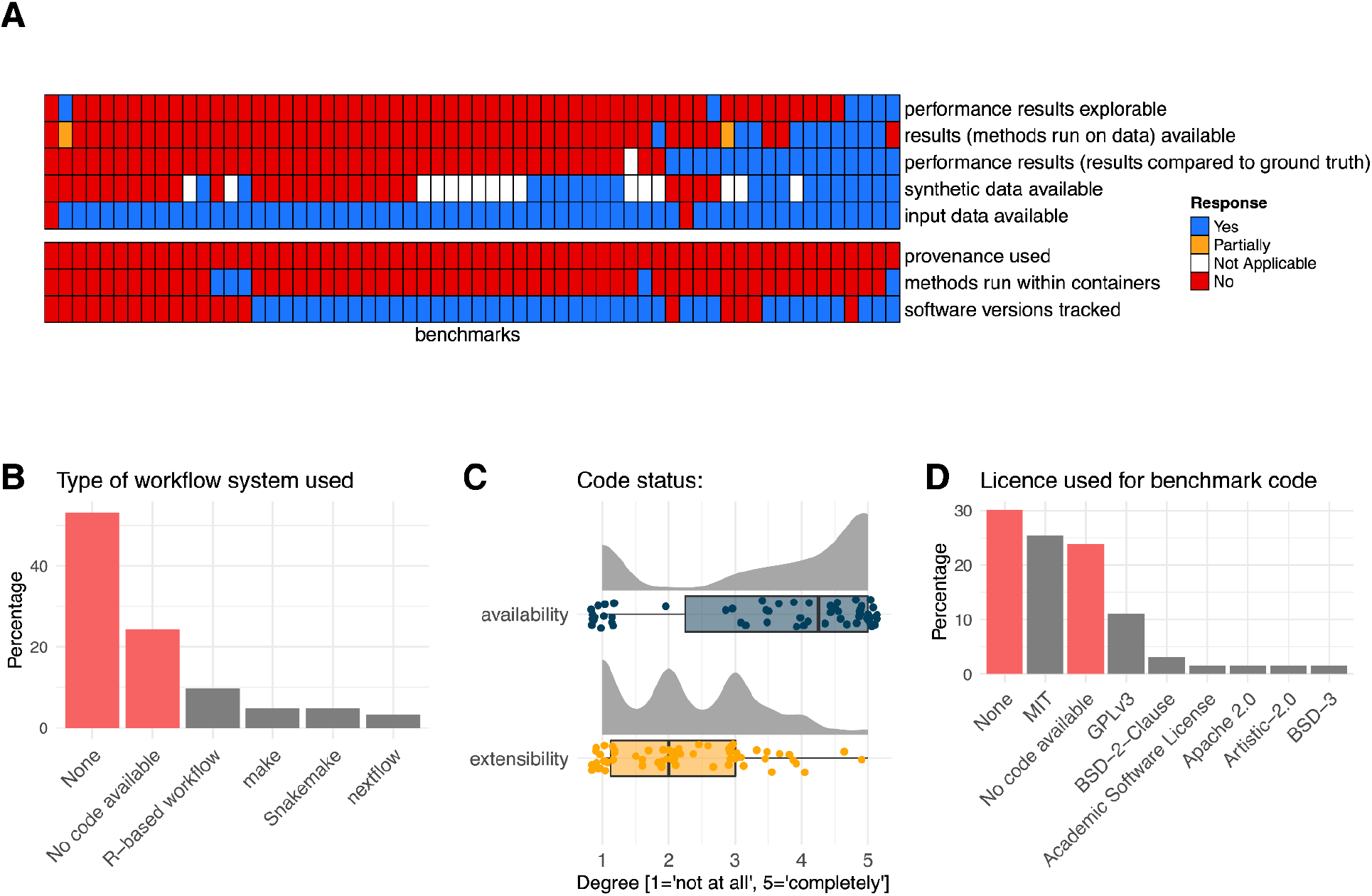
Code / data availability, reproducibility and technical aspects of 62 single-cell method benchmarks. A) Each column of the heatmap represents a benchmark study and each row represents a factual question; responses are represented by colours (*Yes*: blue; *Partially:* orange; *Not Applicable:* white; *No*: red). *Not Applicable* corresponds to benchmarks that did not use simulated data (*synthetic data is available* row) and to a benchmark that evaluated secondary measures only (*performance results available* row); B) type of workflow system used (benchmarks with no workflow used or no code available are represented in red, otherwise grey); C) reviewer’s opinions on the availability and extensibility of benchmarking code; D) licence specification across benchmarking studies (benchmarks without licences or no code available are represented in red, otherwise grey).

We next scored the *degree* to which code is available on a scale of 1 to 5 (Figure 2C) where 1 means ‘not at all’ and 5 means ‘completely’. For over 75% of the benchmarks, code was fully or partially available, such as in a GitHub repository, although there are clearly different levels of completeness and description. We also gathered opinions on how extensible the available code is (e.g., how easy it is to incorporate an additional dataset, method or evaluation criterion; see Figure 2C); amongst the 47 studies sharing code, only two studies received a high score for extensibility.

Explicit code licensing is somewhat sporadic for benchmark studies (Figure 2D); of the 47 studies that made code available, 19 (40%) did not specify a licence. This can become an important consideration when re-using (public but licensed) code for building data analysis pipelines or extending benchmarks. Of those benchmarks that specified a licence, the free software MIT and GPLv3 appeared to be the dominant ones.

Taken together, our meta-analysis shows that most benchmarks results are at least in principle reproducible, since code and input data are shared. However, a fully reproducible analysis would also require information about the software environment (e.g., operating system, libraries, packages) or an available container, which are sometimes documented, but often not readily available in the benchmarks that we evaluated. Thus, a significant amount of redundant work would be required to reestablish or extend the vast majority of surveyed benchmarks.

## The future of benchmarking

Benchmarking of omics tools has evolved to become standard practice and remains a crucial part of the development cycle of computational methods. However, the current standard of benchmarking still leaves considerable room for improvement. For example, benchmarks can become out-of-date rapidly, especially in fast-moving subfields such as genomic data analysis. A better standard would involve updating and running benchmarking continuously as new methods emerge and importantly, as better metrics become apparent. The computational biology community has for a long time been liberal about making code available (Deshpande et al. 2021), mirroring the genomics community in making data available (Byrd et al. 2020), but the current standard in benchmarking is often not much more than a code dump, i.e., a minimal record of what was done. Ideally, methodologists get access to not only code and input data, but also full software environments as well as modular workflow systems to orchestrate the running of methods on datasets and evaluations of results against a ground truth. Ultimately, many of the components that form a benchmark (datasets, methods, evaluation metrics) could be potentially re-used (e.g., ground truth datasets re-purposed for a different computational task). What is lacking right now is a general interoperability between components. Across current benchmarks, there is (typically) no mechanism to add new methods, datasets, or metrics, which is an important consideration, since we want to update and push the state-of-the-art for developers, researchers and consumers of our methods.

Seamless and eternal extensibility or reproducibility can be challenging to establish and maintain indefinitely, especially for single research groups and because software toolchains used in computational biology are constantly in flux. The complete absence of provenance tracking also highlights that the intermediate results of a benchmark are not currently being directly reused. For example, if a developer wanted to evaluate their method within the context of an existing benchmark, they would only have access to the bare minimum (usually just the reference datasets and some code); nonetheless, they are expected to re-run all previous methods essentially reconstructing the benchmark from scratch. This signals an opportunity to make intermediate results available in a systematic way, and use provenance tracking more broadly, whereby the results of methods run on datasets are recorded (as well as metric calculations comparing the results to the ground truth), together with the relevant parameters, such that method developers need only to re-execute relevant new components of a workflow. Reusability and extensibility of benchmarks would not only benefit users but also increase the visibility of published benchmarks, not simply a static and dated view of the field. A related point is software environments, since an ongoing challenge in computational biology is to onboard new methods across diverse computing systems, due to possibly different library requirements (e.g., Python version or Bioconductor release) of each software tool. Here, containerization (Gruening et al. 2018) represents a currently untapped opportunity to systematize this for research pipelines as well as benchmarking studies.

Another area where users (data analysts, method developers) could directly interact with the benchmark data products is in the interpretation of performance. With access to the various assessment criteria, ranking strategies could be established that are tailored to specific use cases and experimental settings, instead of that chosen in advance by the benchmarking team.

Despite the considerable time and effort required to set up a serious benchmark (e.g., collecting and standardising datasets with ground truth, installing and onboarding methods, defining and running metrics), it is still most common to build a benchmark from scratch instead of extending existing benchmarks. To put in context, the single-cell community has developed over 80 methods to infer trajectories and over 40 methods to call differentially expressed genes (Zappia and Theis 2021), but there are seemingly few systems to orchestrate (create, extend, and run) neutral benchmarks. The burden of implementing extensible and continuously updated benchmarks does not necessarily need to be on the side of the benchmarker alone, since most benchmarks employ a similar structure (e.g. datasets, methods, metrics) and could use a general benchmarking orchestration system.

Although here we have focused on stand-alone benchmarks, every method developer is obligated to compare their approach to existing ones and also here a systematic approach has the potential to achieve considerable efficiency. Method developers could use an established (vetted) benchmark to evaluate their method, while introducing considerable time savings, avoiding some of the problems of self assessment (Buchka et al. 2021), and increasing transparency while preserving the value of benchmarks for end users.

Besides having a modern technical implementation to organize benchmarks, it will be important that the community populates the system with high-quality tests of methods and finds strategies to constantly re-monitor the suitability of such tests, in a both systematic and community-informed way. However, several questions remain, such as who is responsible for the cost of computation, how to decide on the composition of high-quality tests of methods, and how to best combine performance metrics to rank methods.

In this article, we identified several bottlenecks in current benchmarking policies, mainly related to reproducibility, extensibility and achieving access to useful content. Taken together, to achieve a more transparent and impactful methods assessment ecosystem, we suggest that systematization of benchmarks through containerization, workflow tools and provenance tracking with access to the full results of a benchmark would be required. A comprehensive list of issues and recommendations for benchmarking identified in this article is available in Table 1, but other guidelines were already covered by others (Mangul, Mosqueiro, et al. 2019; Mangul, Martin, et al. 2019; Weber et al. 2019; Aniba, Poch, and Thompson 2010). Notably, a set of principles to achieve higher quality and sustainability, *FAIRsoft*, was recently developed for research software (del Pico, Gelpi, and Capella-Gutiérrez 2022), coupled with a scoring system scheme that allows a more objective assessment and evaluation of FAIR (Findable, Accessable, Interoperable, Reusable) principles; this strategy could also be applied to benchmarks. Our recommendations also echo a recent call to evaluate computational methods using a common platform (Czarnewski et al. 2022). Several initiatives are in motion here, such as ‘Open Problems in Single-Cell Analysis’ and OpenEBench (Lance et al. 2022; Capella-Gutierrez et al. 2017), and other tools may emerge for achieving this desired interoperability. We believe that if benchmark authors and method developers start embracing these tools, future benchmarks will have an enduring impact in guiding computational methods towards a more structured and comprehensive science.

**Table 1.** List of guidelines discussed in the article and their benefit for the community.

| Scope | Main areas of improvement | Suggestions | Outcomes |
| --- | --- | --- | --- |
| 1. Extensibility | Poor extensibility of benchmarks (addition of new components such as methods or metrics). | Manage code with workflow management systems (see above), improve documentation, organise code to allow addition of new components (increase modularity). | Increased potential for code reuse by method developers and overall research article quality. Reduced effort required for future benchmarks with the same scope. Ultimately, improved comparison of results across studies. |
| 2. Output availability | Intermediate and final benchmarking outputs are often not made public or are not explorable. | Provide (intermediate) outputs in a suitable format as supplementary material or make the available code complete enough to fully regenerate intermediate results. | Easier access to information for readers (specific case-studies). Outputs can be reused for other comparison studies. |
| 3. Parameters | Most evaluated methods are run with default settings. | Evaluate the sensitivity of the methods when parameters need fine-tuning. | Users will be more aware of the critical parameters to set when fine-tuning is necessary. |
| 4. Workflow management and containers | Workflow management systems and containers are scarcely used. | Encourage the training of these tools in scientific workshops and undergraduate courses. | Improved reproducibility of benchmarks and increased chance that they will be reused or extended, thus improving their visibility. |
| 5. Code licences | Licences are seldom defined, making any code reuse unclear. | Define a licence based on the scope of the research and potential commercial distribution. | Increased reuse of code and decreased chance of (un)intentional misuses of intellectual property. |

## Methods

To perform the survey, we first gathered a list of published benchmarks in the field of single-cell data analysis. The list was crowdsourced by the project team using search engines, literature searches, our own benchmarks, as well as a call on Twitter. Selected benchmarks had to have at least a preprint published and should not showcase a new computational method. 62 benchmarks covering 19 analysis tasks were gathered and used for the rest of the analysis (see Additional file 1 for the list of studies).

A questionnaire was designed before the collection of data to evaluate different aspects of the benchmarks, grouped into ‘General information’, ‘Code/ data availability’ and ‘Technical aspects’ (see Additional file 2). The latter two categories were merged into ‘Code, data availability and technical aspects’ in the manuscript for more clarity and because they were regrouping information from a similar aspect. The questions covered both factual and opinion assessments. The factual questions could be numerical (“Number of methods evaluated”), categorical (“Type of workflow used”) or semi-open (e.g.: answers to “What secondary measures were assessed?” could be of “Code quality”, “Installation” or a free answer to specify). All reviewers are students, postdocs or group leaders actively involved in computational biology methods development. Several of the reviewed benchmarks were performed by our research group; however, to ensure neutrality, we allocated reviewers to be someone that was not directly involved. Except for this constraint, the reviewers could freely assign themselves to any benchmarks that corresponded to their own domain of expertise or interest. Each benchmark was reviewed at least twice by two reviewers (see Additional file 3 for original responses). Some questions received a high level of concordance among the two reviews (example, “Type of workflow system used” and “Whether input data used by the methods is available”) while some received a lower level of concordance (“Degree to which code to re-run benchmark is extensible”, “What secondary measures were assessed?”). To resolve discrepancies, we employed a third reviewer to harmonise one or several questions across all benchmarks. Most discrepancies came about from unanswered questions, adding text to numerical answers, or simple errors (e.g., missing that a code repository was made available). A small number of discrepancies arose from underestimations due to relevant informations being missed during one of the reviews (e.g., “Degree to which authors are neutral”, where some reviewers missed an author’s contribution to one of the evaluated methods), while other harmonizations required a re-analysis of the answers and a discussion with individual reviewers (e.g., “Degree to which code to re-run benchmarks is extendible”). The analysis presented in this manuscript was performed on the harmonised answers.

## Contributions

All authors contributed to the review of at least one benchmark. MDR designed the analysis and the questionnaire, with input from all authors. AS, AL, CS, IM, PLG and MDR harmonised the reviews from the questionnaire, analysed the data and contributed to the writing of the manuscript.

## Additional files

Additional file 1 - list of benchmarks with consolidate survey answers (CSV)

Additional file 2 - review form (CSV)

Additional file 3 - unedited (anonymized) survey responses (TXT)

## Code and data availability statement

The code to organize, consolidate, monitor and analyze the benchmark reviews is available from https://github.com/markrobinsonuzh/sc_benchmark_metaanalysis and a snapshot is posted at zenodo (DOI: 10.5281/zenodo.7097767). The additional files mentioned above are available at zenodo (DOI: 10.5281/zenodo.7733753).

## Supplementary Figure

**Supplementary Figure 1.**
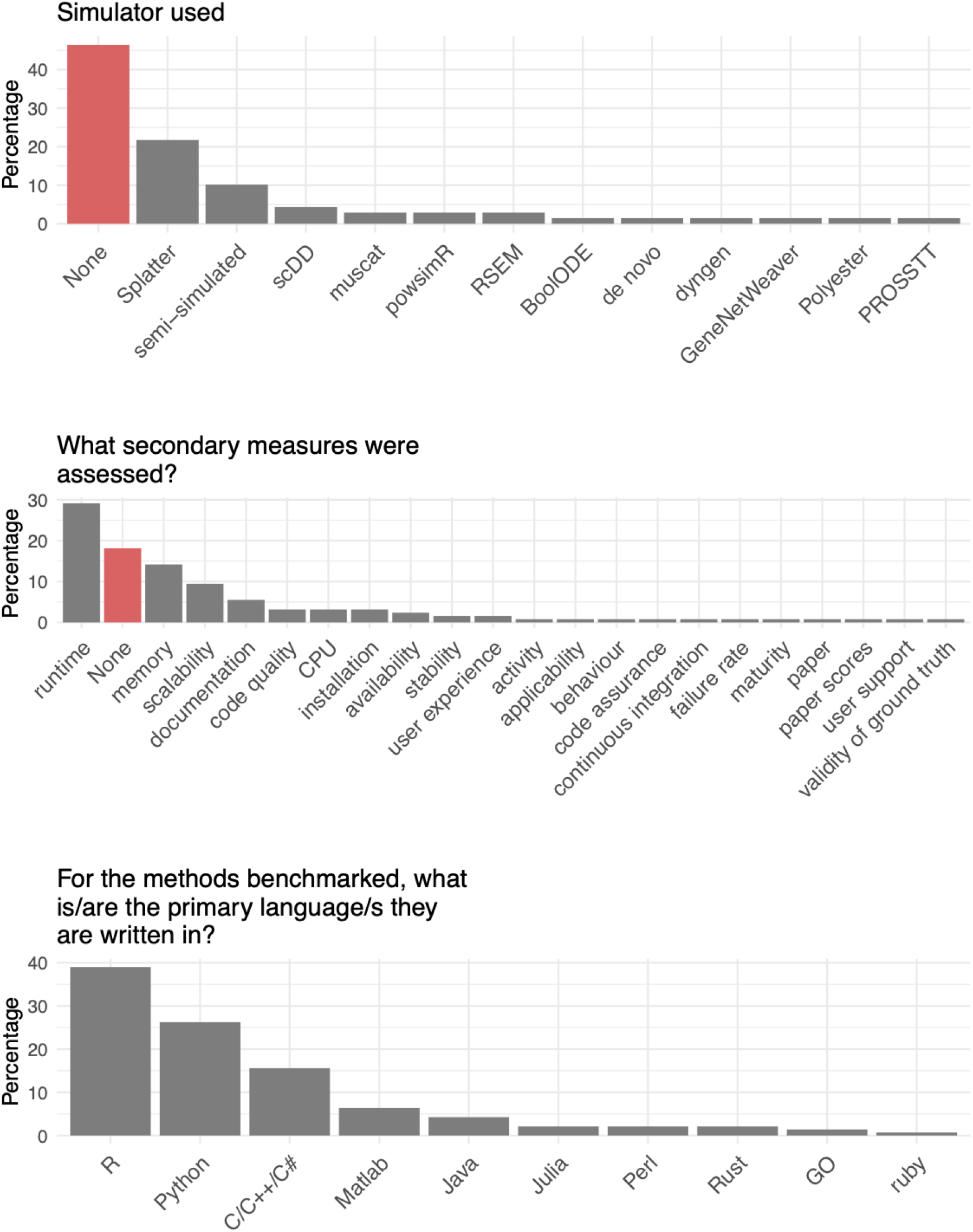
Auxiliary summaries from the survey responses. Simulator used, secondary measures assessed, language of tools assessed.

